# Genome streamlining to improve performance of a fast-growing cyanobacterium *Synechococcus elongatus* UTEX 2973

**DOI:** 10.1101/2024.01.16.575707

**Authors:** Annesha Sengupta, Anindita Bandyopadhyay, Debolina Sarkar, John I. Hendry, Max G. Schubert, Deng Liu, George M. Church, Costas D. Maranas, Himadri B. Pakrasi

## Abstract

Cyanobacteria are photosynthetic organisms that have garnered significant recognition as potential hosts for sustainable bioproduction. However, their complex regulatory networks pose significant challenges to major metabolic engineering efforts, thereby limiting their feasibility as production hosts. Genome streamlining has been demonstrated to be a successful approach for improving productivity and fitness in heterotrophs but is yet to be explored to its full potential in phototrophs. Here we present the systematic reduction of the genome of the cyanobacterium exhibiting the fastest exponential growth, *Synechococcus elongatus* UTEX 2973. This work, the first of its kind in a photoautotroph, involved an iterative process using state-of-the-art genome-editing technology guided by experimental analysis and computational tools. CRISPR/Cas3 enabled large, progressive deletions of predicted dispensable regions and aided in the identification of essential genes. The large deletions were combined to obtain a strain with 55 kb genome reduction. The strains with streamlined genome showed improvement in growth (up to 23%) and productivity (by 22.7%) as compared to the WT. This streamlining strategy not only has the potential to develop cyanobacterial strains with improved growth and productivity traits but can also facilitate a better understanding of their genome to phenome relationships.

**Importance:** Genome streamlining is an evolutionary strategy used by natural living systems to dispense unnecessary genes from their genome as a mechanism to adapt and evolve. While this strategy has been successfully borrowed to develop synthetic heterotrophic microbial systems with desired phenotype, it has not been extensively explored in photoautotrophs. Genome streamlining strategy incorporates both computational predictions to identify the dispensable regions and experimental validation using genome editing tool and in this study we have employed a modified strategy with the goal to minimize the genome size to an extent that allows optimal cellular fitness under specified conditions. Our strategy has explored a novel genome-editing tool in photoautotrophs which, unlike other existing tools, enables large, spontaneous optimal deletions from the genome. Our findings demonstrate the effectiveness of this modified strategy in obtaining strains with streamlined genome, exhibiting improved fitness and productivity.

## Introduction

Cyanobacteria are the most ancient and abundant oxygenic photosynthetic organisms that are largely responsible for the Earth’s viable environment (1). These photosynthetic prokaryotes have been identified as potential platforms for sustainable carbon-neutral bioproduction because of their unique ability to harvest sunlight as their sole energy source for converting greenhouse gases (carbon dioxide) into value-added chemicals. This bioprocess in theory is also economically sustainable as it enables free and ubiquitous substrates to enter the bioeconomy. In comparison to other photoautotrophs, cyanobacteria have several advantages and therefore, efforts are ongoing to understand and develop the cyanobacterial platform as sustainable bio-factories (2-5). However, relatively slower growth and limited knowledge of their genomic traits as compared to heterotrophs such as *E. coli* or yeast have restricted progress in this direction. Though the recent isolation of a few fast-growing strains have made cyanobacterial-based bioproduction more compelling and tractable than ever, concerted efforts are needed to get their productivity at par with their heterotrophic counterparts (6-10). Although most commonly studied cyanobacterial genomes are generally smaller in size than that of *E. coli*, they exhibit cryptic metabolic and regulatory features, owed likely to their photosynthetic lifestyle, unique evolutionary history, and adaptation to various unfavorable environmental conditions(11).

The main aim of this study was to test the feasibility of employing the genome reduction strategy as a means to shed excess biological complexity and simplify the genome of a fast-growing cyanobacterium without compromising its desirable traits. The clade *Synechococcus elongatus* hosts all of the fast-growing cyanobacterial strains identified to date (8) and a genome minimization approach will be beneficial for unravelling the genome level function and the overall metabolism of these strains which in turn will aid their development into cell factories. The goal is not to obtain a truly minimum genome for a photosynthetic organism, but rather identify and remove genes dispensable under bioproduction-relevant conditions (high light and CO_2_) without compromising growth and productivity.

Genome streamlining is a natural evolutionary process of eliminating non-beneficial genes, since a smaller genome reduces the metabolic burden on the cell and improves fitness (12-14). As an engineering strategy, it’s been successfully employed in model heterotrophs leading to improved fitness and performance(15-18). A genome reduction of 25% in *E. coli* led to a 1.6-fold improvement in growth rate as well as improved recombinant protein production (19, 20). Similarly, genome streamlining in a *Pseudomonas* strain resulted in several appealing traits such as faster growth, increased biomass production, enhanced plasmid stability, and overall a more efficient energy metabolism (21, 22). These studies indicate that a chassis strain with a streamlined genome avoids the unnecessary burden of replicating and expressing genetic elements that are not useful under production conditions. Reducing this unnecessary genetic burden may ensure more cellular resources are available for expression of heterologous pathways. By removing genes of unknown function, it also creates a chassis organism that is more fully-understood, more amenable towards genetic engineering and synthetic biology efforts (16). Despite the favorable outcomes of genome minimization in heterotrophs (20-23), limited efforts have been made to implement such strategies in phototrophs. So far two reports explore genome streamlining. Removing ∼2% (118 kb) of the genome of *Anabaena* PCC 7120 has been demonstrated using the CRISPR-Cas12a system for targeted deletions, but this study did not explore the effects of these edits on strain performance (24). Deletion of several large fragments of DNA from the genome of *Synechococcus elongatus* PCC 7942 has been performed, producing a septuple mutant with approximately 3.8% of its genome removed. These mutants were further studied to understand the changes in the transcriptomics profile of the cells resulting from the deletions. The CRISPR-Cas12a editing tool was used to obtain the targeted, specified and markerless deletions in this study (25). Though Cas12a is a dynamic and versatile tool, it does not allow the flexibility to explore and identify unknown stretches of dispensable and indispensable regions in the genome of a strain. Recently, a novel Class I multi-Cas protein was commissioned for large deletions in heterotrophs. The dual helicase and exonuclease activity of Cas3 enabled large simultaneous bi-directional deletions, without the necessity of a repair template (26). Recently in one of our studies, we have successfully commissioned an inducible CRISPR Cas3 system in *Synechococcus elongatus* UTEX 2973 to truncate the light-harvesting antenna structure for maximizing fitness and productivity under specified condition (27). However, this tool has not been explored for genome streamlining in cyanobacteria. Therefore, it is of interest to investigate this Cas system and exploit its beneficial features for genome minimization of photoautotrophs.

In this study we investigate the effect of systematic reduction of dispensable regions from the genome of *Synechococcus* 2973, the fastest growing, high light thriving cyanobacterium that exhibits high sucrose production titer (6, 10). We first identified five large stretches of dispensable genomic regions in this strain using the MinGenome algorithm (28) and then commissioned the novel CRISPR-exonuclease system, CRISPR-Cas3 to achieve flexible progressive large deletions. CRISPR-Cas3 besides deleting large regions, enabled identification of non-dispensable genes in *Synechococcus* 2973 which otherwise were predicted as dispensable. We successfully deleted the optimal stretch of two of the five dispensable regions identified by our *in silico* analysis. The two large deletions were combined to obtain a strain with genome reduction of 55 kb. The strains with reduced genome showed improved growth and sucrose productivity. This proof-of concept study (Fig. 1) demonstrates that systematic minimization of cyanobacterial genomes has the potential to develop these organisms as super-strains that might hold the potential to boost a carbon-neutral bio-economy and mitigates climate issues.

**Fig. 1:**
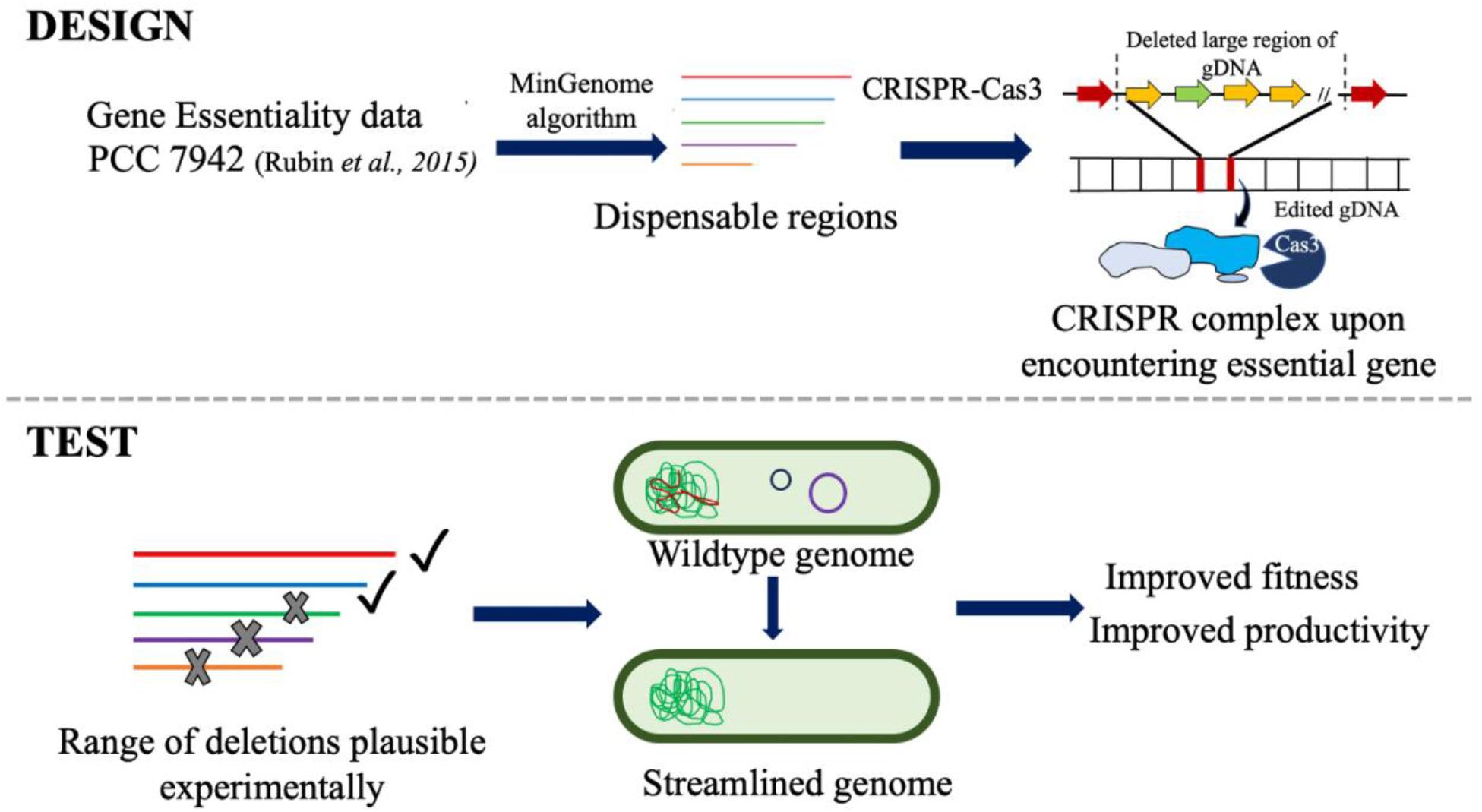
The schematic showing the strategy used for systematic genome streamlining in cyanobacteria. A combinatorial approach integrating computational (design) and experimental (test) tools to first identify the dispensable regions in *Synechococcus elongatus* UTEX 2973 and further use novel CRISPR tool to validate the prediction and create a strain with improved fitness and productivity.

## Result

### In silico prediction of probable dispensable regions from the genome of Synechococcus 2973

The first step towards streamlining the genome of *Synechococcus* 2973 was to identify the dispensable genomic regions (Fig. 1). Since *Synechococcus* 2973 shares >99% genomic identity with the model strain *Synechococcus* 7942, the list of essential genes already available for *Synechococcus* 7942 (29) was borrowed and mapped to *Synechococcus* 2973 using a bidirectional protein blast with a stringent e-value cut-off of 10^−10^ to avoid spurious hits. The MinGenome algorithm (28) was employed to predict large dispensable regions with three criteria: (1) The growth rate of the strain is fixed at the maximum, (2) Essential genes mapped from *Synechococcus* 7942 cannot be deleted, and (3) Find the longest stretch first and iterate to find the consecutive ones. This strategy predicts five large regions of the genome can be dispensed without affecting the strain fitness (Table 1). The list of gene annotations of each cluster are provided in Table S1-S5.

**Table 1:** A list of large gene clusters that are predicted to be unessential for growth of *Synechococcus* 2973. The predicted dispensable region in between the start site of the first gene of the cluster and the start site of the end gene of the cluster.

| Rank | Start | End | Length (bp) |
| --- | --- | --- | --- |
| 1 | M744_12940 | M744_13140 | 33,952 |
| 2 | M744_10800 | M744_10895 | 26,399 |
| 3 | M744_02780 | M744_02900 | 25,699 |
| 4 | M744_12500 | M744_12615 | 22,544 |
| 5 | M744_05410 | M744_05555 | 22,235 |

### CRISPR Cas3-mediated genome editing enabled deletion of large genomic regions

A Rhamnose-Theophylline inducible Class I CRISPR system involving multi-Cas proteins was employed to streamline the genome of *Synechococcus* 2973 (Fig. 2a). In this CRISPR system, Cas3 is the key component exhibiting both helicase and nuclease activity, enabling large bidirectional deletions. The other Cas proteins (Cas5, 7, and 8) help in maintaining the fork structure required for these long deletions (26, 27). Unlike Cas12a-mediated editing where the length of deletion is pre-determined based on existing knowledge of the region of interest (designed repair template), the Cas3 system allows spontaneous deletion of DNA stretches on either side of the targeted region, as long as the deletions are not detrimental for the strain. Therefore, the Cas3 CRISPR system has the potential to not only identify previously unknown dispensable regions in the genome but also uncover essential genes within a stretch of DNA that has been computationally determined as dispensable, thereby averting any adverse effects on the strain from large scale deletion experiments.

**Fig. 2:**
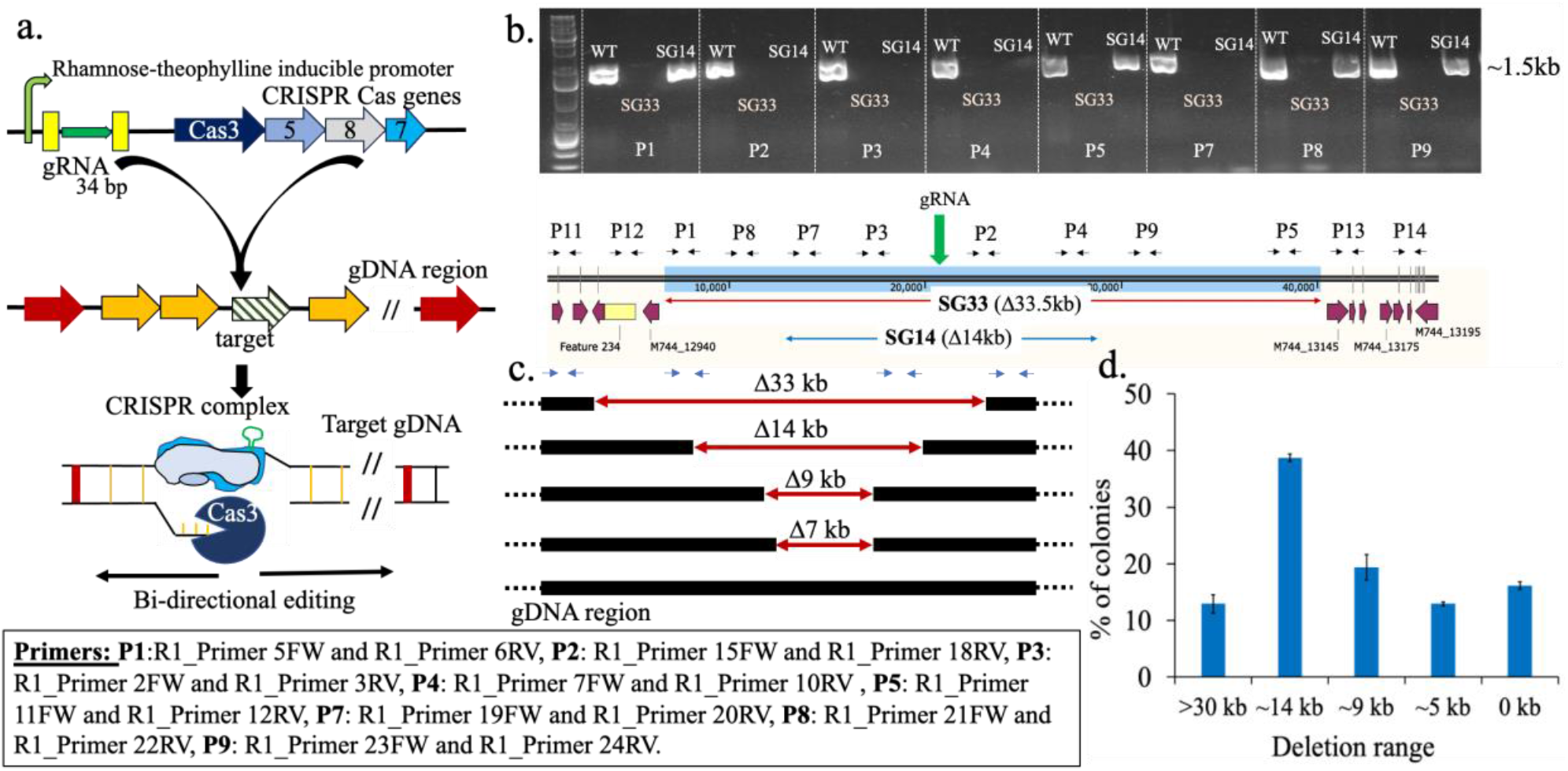
Developing CRISPR-Cas3 system for spontaneous large deletions in cyanobacteria. (a) The schematic showing the functioning of the CRISPR-Cas3 system. (b) The result from Tiling PCR (gel picture) using different primer sets (P1-9) showing the absence or presence of 1.5 kb band for two colonies (SG33 and SG20). The absence of band indicate the region is deleted from the strain as compared to WT. (c) The length of deletions obtained when CRISPR system was targeted at the R1 region. (d) Frequency of occurrence of colonies of a particular length of deletion. The error bars represent the variation observed between the difference in the number of colonies obtained for a particular length of deletions between three separate experiments.

In our previous study, as a proof of concept, the inducible CRISPR-Cas3 system was tested to delete dispensable genes from the genomic cluster encoding the light-harvesting antenna complex of *Synechococcus* 2973. Interestingly, every deletion attempt led to a strain with a reduction of 4 kb. Investigation of the region beyond 4kb revealed that the genes immediately flanking the deletion on either side were designated as essential as per the gene essentiality data (27). These experiments revealed the potential of the Cas3 tool in identifying essential genes in a stretch of DNA while simultaneously identifying dispensable regions. Thus our next goal was to test the feasibility of using the Cas3 entourage for deletions of large genomic segments from cyanobacterial genomes, analogous to what has been achieved in heterotrophs. In this work, a strategy similar to a heterotrophic study (27) was implemented to target all the 5 regions predicted as dispensable by our *in silico* analysis (Table 1). We first commissioned CRISPR-Cas3 for the largest stretch identified, R1 (Table 1), in order to optimize the system for large deletion in *Synechococcus* 2973 (Fig. 2). A 34 bp long gRNA was designed for R1 that targeted the gene M744_13025 (which is approximately at the midpoint of the predicted region). Around 30 colonies out of 1000 were initially screened for deletions by using tiling PCR with 3 sets of primers (P1, P3 and P4) and around 20% of the colonies showed no deletions (Fig. S1). The probable position of the primers initially chosen are shown in Fig. 2b. A more extensive tiling PCR with multiple primers helped identify the range of deletions in different colonies, such as SG33 and SG14 (Fig. 2b). Similar analysis of all the rest 80% positive colonies indicated a wide range of deletions from 0 to >30kb (Fig. 2c). A frequency analysis showed around 40% of the deletions were in the range of 14-15kb and ∼10% colonies showed >30 kb deletion (Fig 2d). This demonstrated that the CRISPR-Cas3 tool has the potential to execute a large range of deletions in cyanobacteria, generating a library of random deletions.

This tested inducible CRISPR-Cas3 system was then used to create strains with streamlined genome by targeting the predicted regions (Table 1). Fig 3a briefly shows the workflow for generating these deletion mutants using this inducible CRISPR-Cas3 system. CRISPR-mediated targeting of the R1 region gave rise to a markerless strain 2973Δ33.5kb showing a deletion of 33.5 kb region (hereafter mentioned as SG33) (Fig. 3). The extent of deletion in the strain was initially confirmed by tiling PCR (Fig. 3b), and then the total length of deletion was determined by confirmation PCR using set of primers that were previously determined based on the tiling PCR results (Fig. 3b). Using primers flanking the 40kb targeted region, a band of ∼7kb was obtained in the SG33 strain, indicating a large deletion of ∼33kb (Fig.3c,d). Finally, whole genome sequencing confirmed the range of deletion to be 33.5kb (Fig. 3e). The WGS also confirmed no additional mutations in the SG33 strain.

**Fig. 3:**
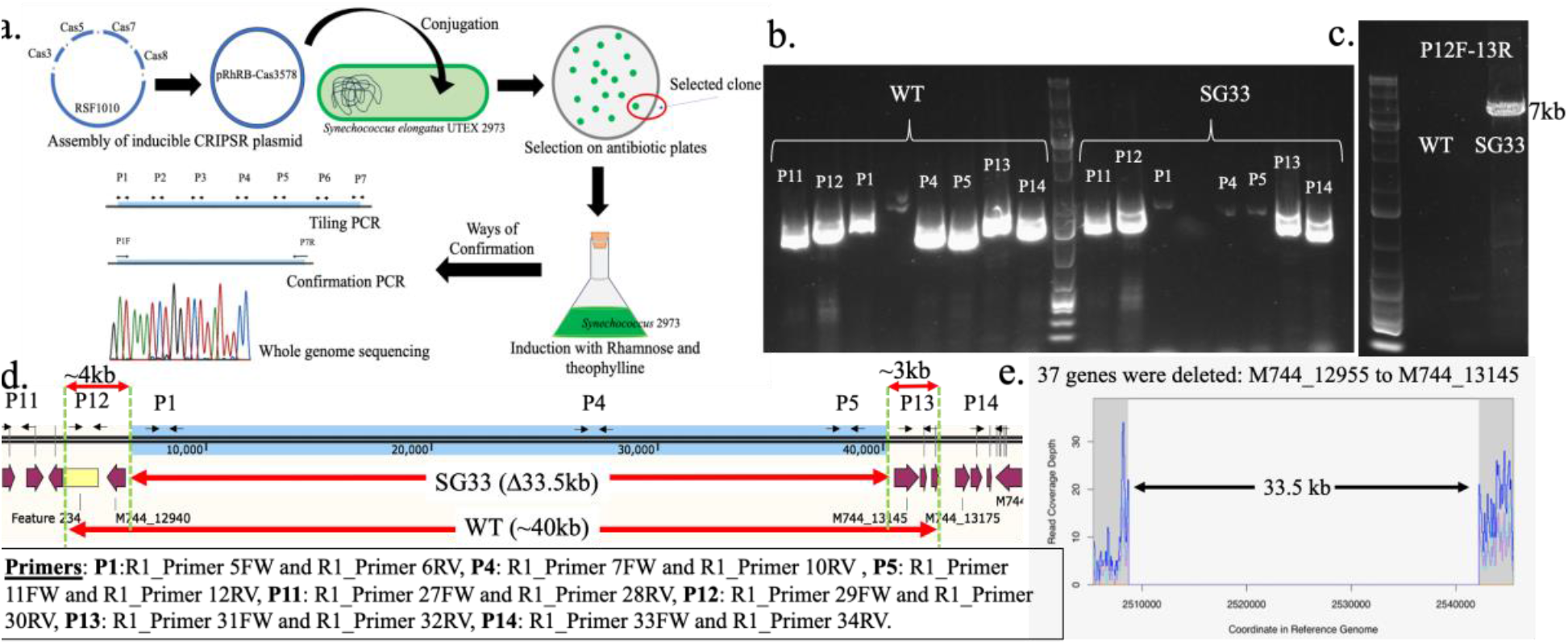
Creating the SG33 strain using CRISPR-Cas3 system. (a) The workflow for obtaining the markerless strain with minimized genome. (b) The gel picture showing the absence or presence of 15 kb band when amplified with different primer sets in a tiling PCR to identify the deleted regions. (c) The Confirmation PCR showing the presence of 7kb band in SG33 strain as opposed to the 40 kb band. No band in WT as the 40 kb band is difficult to obtain. (d) The schematic showing the R1 region and the range of deletions and the region of primers binding. (e) The whole genome sequencing of the SG33 strain showed a clean and segregated deletion.

Similar workflow was applied for the R2 region which led to its partial deletion. Though this region was predicted to be ∼26.4 kb long (Table 1), experimentally the largest deletion obtained from was of 19.7 kb, giving rise to the strain 2973Δ19.7 kb (hereafter mentioned as SG20) (Fig 4). While the first two regions, R1 and R2 could be deleted (Fig. S2), the other identified regions (R3, R4, R5) could not, despite several attempts with different gRNA and PAM sequences (Fig. S3). We then attempted to delete these regions using the CRISPR-Cas12a system with 2 gRNAs (24, 30), but remained unsuccessful as well (data not shown). This suggested that the regions predicted as dispensable based on S7942 essentiality data are probably not accurate enough and indicated the need for *Synechococcus* 2973-specific essentiality data.

**Fig. 4:**
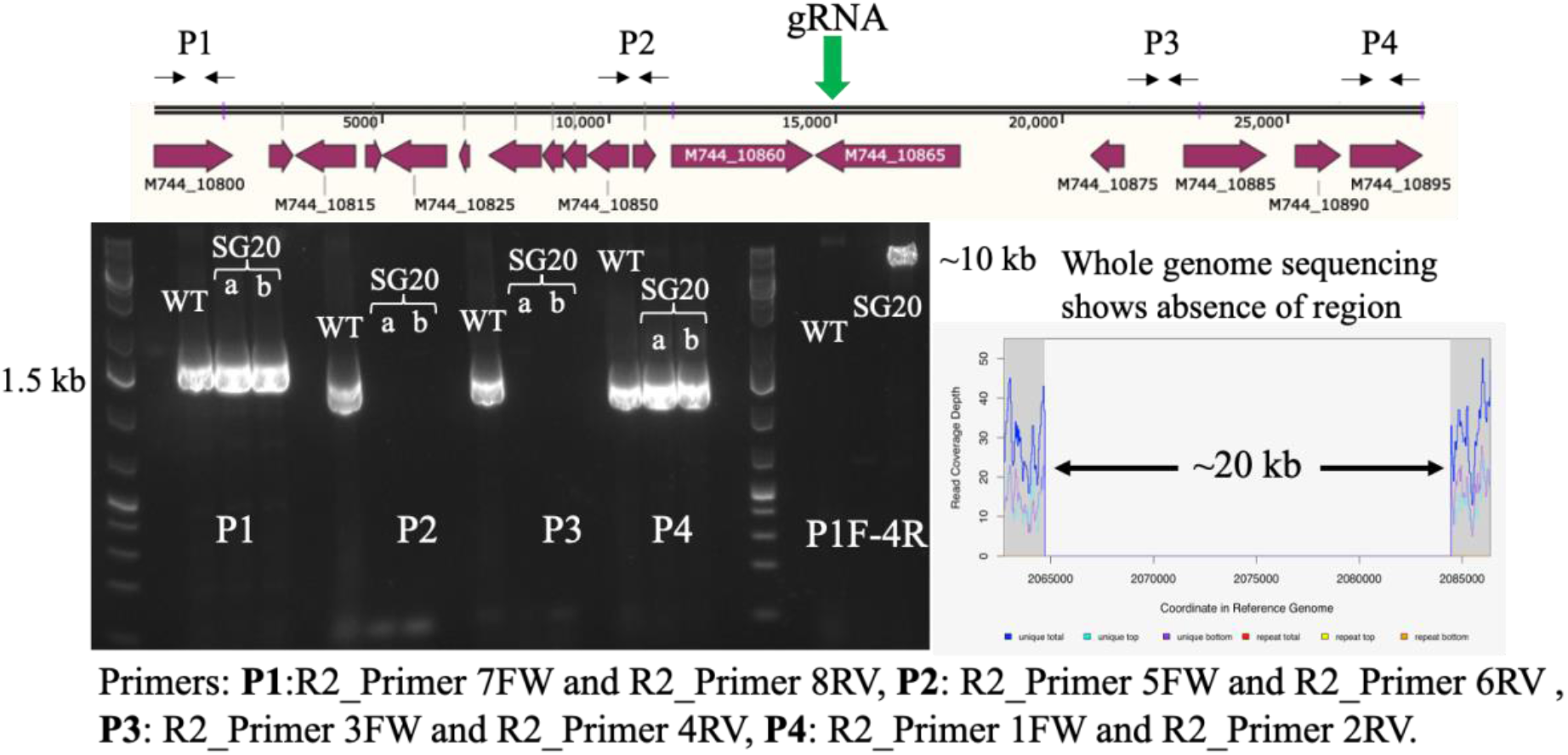
Generation of the SG20 strain. The schematic showing the R2 region and the primer binding sites. The gel picture is of the tiling PCR performed using the different primer sets (P1-P4) and the confirmation gel using the primers P1Forward and P4 Reverse. A 10 kb band is visible instead of the 30kb band in SG20 as compared to the WT. No band is visible as 30kb is difficult to amplify and visualize. A snap of the whole genome sequencing chromatogram shows the absence of reads around the R2 region. Two different colonies with deletion were tested (a, b) as shown in the gel picture.

We then proceeded to combine the two large deletions (SG33 and SG20) to obtain a single strain with a large reduction in the genome. Initially, SG33, the strain with the largest deletion, was used as the base strain to tier the next largest deletion. However, the conjugation efficiency of this strain was very poor [1000 fold lower than the WT or the SG20 strain (data not shown)]. Therefore, we decided to use the SG20 strain as the base strain into which the plasmid carrying the Cas3 machinery for R1 deletion was introduced. Whole Genome Sequencing of one of the transformant colonies showed a total deletion of 55kb in this strain (henceforth referred to as SG55) (2973Δ19.7kbΔ34.3kb). Unlike SG33, where 33.5kb was deleted upon targeting R1, a 34.3 kb region could be deleted when the same region was targeted in the SG20 strain. The above results confirmed the efficacy of the CRISPR-Cas3 genome editing tool in creating spontaneous and large deletions in cyanobacteria.

### Minimization of genome showed improved growth and productivity

The strains with streamlined genome were characterized to determine the fitness and photosynthetic productivity. Since our primary goal was to develop a reduced genome cyanobacterial host, that can thrive under high light and high CO_2_, all studies were performed under high light (1500μmoles.m^-2^.s^-1^) and high CO_2_ (1% CO_2_ bubbling) conditions. A comparative analysis of growth rates revealed an increase of 23%, in the SG33 strain compared to the WT. In contrast, a growth rate increase of only 5% was observed in SG20, while SG55 showed an improvement of 9% (Fig. 5a, Table S6). However, this enhancement in growth rate was not evident under conditions where carbon was a limiting factor (Fig. S4). Interestingly, the SG55 strain showed a 10 fold higher transformation efficiency with RSF1010 plasmids compared to the WT. Analysis of the genome sequence of this strain indicated that this increase in efficiency could likelybe due to a mutation it acquired in SeAgo, an argonaute protein known to reduce the efficiency of RSF1010 plasmid transformation (31).

**Fig. 5:**
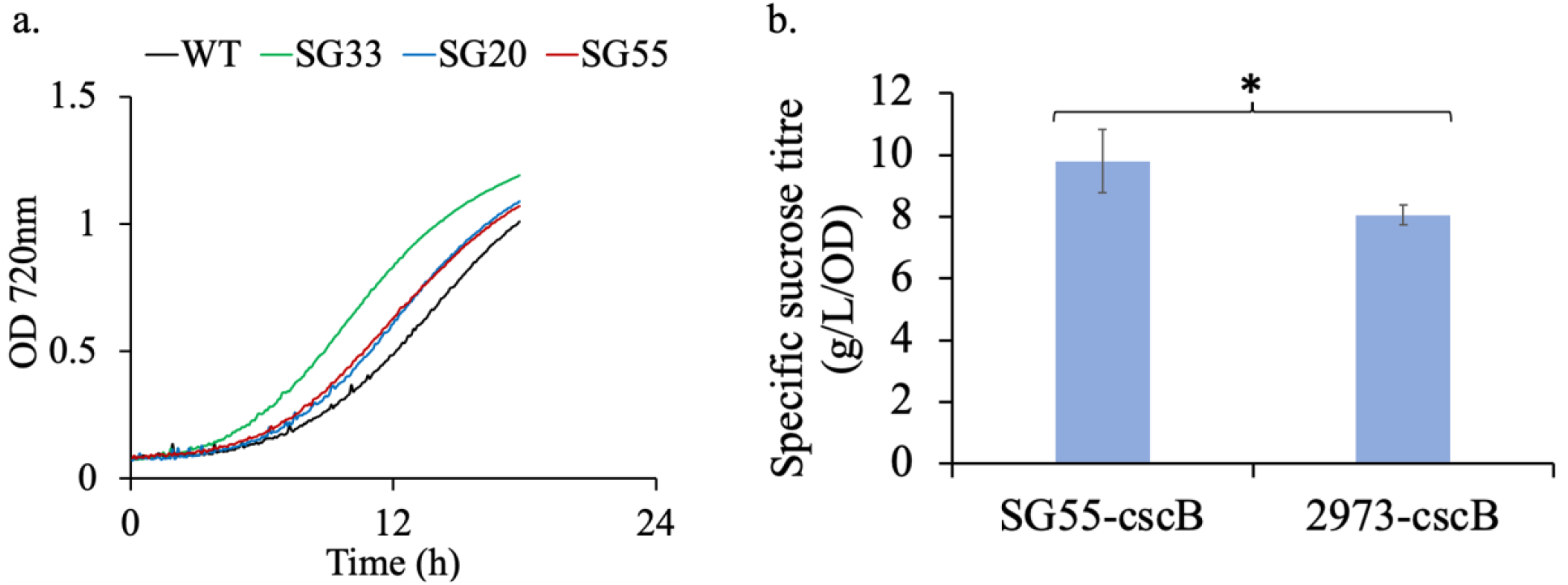
Characterizing the engineered strains with minimized genome. (a) Growth comparison of WT, SG33, SG20 and SG55 under high light and high CO_2_. Experiments were performed in triplicates (n=3) and the graphs are representative growth profile. (b) Sucrose productivity of SG55 as compared to WT. Experiments were performed in triplicates (n=3) and the error bars correspond to standard deviations from at least 3 biological replicates. Asterisk (*) denote statistically different values of μ (p < 0.05).

In addition to the large chromosomal deletions, the two endogenous plasmids pANL (∼50kb) and pANS (∼7kb) were also deleted from the WT strain. Since these deletions did not show any added growth advantages (Fig. S5), these deletions were not tiered into the SG55 strain. Analysis of the photosynthetic efficiency of the strains with streamlined genomes revealed no difference in quantum yield (Fv/Fm 0.35±0.08) as compared to the WT, indicating no adverse impact of the deletions on key physiological processes. Thus, the streamlining strategy employed in this study reduced the metabolic burden in the WT *Synechococcus* 2973 strain resulting in faster growth but did not disrupt its basic cellular functions.

The final strain with streamlined genome (Δ55kb) and improved phenotypic characteristics was engineered to secrete sucrose by integrating *cscB*, a heterologous sucrose transporter gene (10, 32) into the genome. The engineered SG55 strain (SG55-cscB) showed 22.7% higher sucrose production as compared to the strain expressing the transporter alone (10) (Fig 5b). Our findings demonstrate that genome streamlining in cyanobacteria can be a promising strategy for improving both productivity and fitness, provided the correct genes are dispensed.

### Analysis of the deleted regions revealed a large number of hypothetical genes

Streamlining of genome is an effective strategy not only for developing a strain as a production host with beneficial features but analysis of the dispensable genes might also help deduce the role of certain genetic elements. In our analysis, we observed that in the predicted dispensable regions >50% of genes are annotated as hypothetical and majority of these hypothetical genes are in R1 and R2 (Table S1-S5), therefore deciphering the underlying reason for the observed phenotype of the deleted strain was not possible with certainty.

Since in this study the gene essentiality data, was borrowed from *Synechococcus* 7942 (29), there are some discrepancies between the experimentally obtained deletions and predicted dispensability data. To exemplify, in the R1 region there are 37 genes (M744_12940-M744_13135) and the genes flanking this region have been categorized as essential. Conversely, our sequencing results for the SG33 strain which has the R1 deletion, showed the deletion of M744_13140, a gene annotated as essential. Though for non-model strains, strain-specific essentiality data is important, the use of the novel Cas3 system provided the flexibility to dispense the optimal length of regions from the genome for improved fitness. Therefore, the genome streamlining with the CRISPR-Cas3 system is an advantageous strategy and a more high throughput streamlining effort might help create a library of strains exhibiting varied phenotypes.

## Discussion

Genome streamlining is a synthetic biology approach which allows strategic reduction of the genome for attaining a desirable strain phenotype and this approach has been demonstrated to be successful in heterotrophs. There are two approaches of genome streamlining, bottom-up and top-down (33). Though most minimized genome studies have employed a bottom-up strategy to create truly minimal genome from scratch (16, 34), this strategy can pose several challenges as it demands a vivid and thorough knowledgebase of all biological processes and interactions and requires efficient synthetic DNA synthesis and assembly tools. Moreover, cyanobacteria being a polyploid organism exponentially enhances the challenge of introducing and maintaining the synthetic chromosome. The other approach is top-down, which involves systemic streamlining of the existing genome based on prior information regarding the core essential genes. Most genome streamlining efforts have relied on traditional and more recently, the CRISPR-Cas12a/9-mediated genome editing tools (20, 21, 24, 25). For strains where the essentiality information is available, the strategy to delete fixed, specified regions of the genome is advantageous as it leaves less room for casualties such as drastic loss of fitness. However, even with prior knowledge, the experimental outcome might not correlate with *in silico* predictions. Like, in *E. coli* MG1655 a reduction of 29.7% exhibited severely impaired phenotype, while 7% reduction showed no retarded phenotype (20). Therefore, for known and more importantly for newly discovered strains, the spontaneity or randomness in the extent of deletion might be the key to obtain a strain with enhanced features, and CRISPR-Cas3 mediated genome editing has the potential (26, 27). The RNA-guided Cas3 protein has dual helicase-nuclease activity which allows large, progressive, random deletion of genomic region unless encountered with an essential gene. CRISPR-Cas3 has been demonstrated to be effective in heterotrophs, such as *Pseudomonas aeruginosa*, where deletion of genomic regions as large as 424 kb with a mean of 92.9 kb and median of 58.2 kb deletion (26) was observed. Therefore, its use in photoautotrophs for streamlining and other large scale editing purposes is worth exploring.

In this study, we focused on developing a genome streamlining strategy for a non-model cyanobacterium *Synechococcus* 2973 to obtain a strain with reduced genome and improved fitness. *Synechococcus* 2973 is the fastest growing cyanobacterium known so far with a doubling time comparable to heterotroph model strains such as yeast (6, 35). This organism has the potential to be developed as the next-generation production host, however, the complexity of the strain poses challenges for major engineering efforts. We attempted to minimize the metabolic burden on this strain by first identifying dispensable genomic regions using the MinGenome algorithm (28) and then removing them under bioproduction-relevant conditions without compromising growth and productivity (Fig. 1). This iterative integrated approach led to the creation of engineered *Synechococcus* 2973 strains with minimized genomes exhibiting significant growth advantage (Fig 5a). Our results indicate that the extent of growth advantage is not dependent on the extent of genome reduction but on the set of deleted genes. Our analysis revealed that some genes in R1 are predicted as phage-associated proteins. Though SG55 strain has the largest range of deletion (55kb), the growth improvement is more in SG33 (33.5kb deletion) and this might be due to the deletion of prophage-like genes. A previous study in *Vibrio natriegens* revealed that deletion of prophage-containing genomic regions is an effective engineering strategy for improving growth (36).

Since a majority of the genes are annotated as hypothetical, further analysis to decipher the genetic features responsible for the phenotype was not possible. We tested the effect of minimization on productivity by engineering SG55 strain for sucrose overproduction and observed a 22.7% improvement in sucrose titer (Fig. 5b) as compared to sucrose producing WT (10). This strategy of streamlining the genome for improved growth and phenotype might come at the cost of robustness under non-controlled conditions such as outdoor like conditions (carbon limited). Under carbon limited condition, the engineered strains failed to show the improved phenotype (Fig. S4). This decreased fitness in conditions mimicking natural environment (limited carbon available) might be a boon as it lessens the risk of their release and the chance to overtake the WT populations. However in under elevated CO_2_, these strains might be advantageous to utilize more available carbon.

A more high throughput understanding of the phenotype and genotype relationship would provide a better insight into the complexity of the strains. The stochastic nature of the CRISPR-Cas3 system offers the potential to discern essentiality of genes in the targeted regions and obtain an optimal dispensable stretch while retaining or enhancing strain fitness. This hybrid strategy of combining computational analysis with progressive deletion of genomic regions can generate a library of engineered model as well as non-model photoautotrophic strains with streamlined genomes, thereby paving the path for developing these remarkable green hosts into predictable bio-systems with potential to mitigate climate problems and boost bio-economy.

## Materials and Methods

### Computationally predicting longest dispensable regions

To identify regions in the *Synechococcus* 2973 genome that can be deleted without affecting organism growth, we used a genome reduction algorithm called minGenome (28). minGenome uses a mixed-integer linear programming based approach to systematically identify (in descending order) all contiguous dispensable regions. First, essential genes were identified using sequence homology with *Synechococcus* 7942 using a bidirectional protein BLAST. These can then be flagged and their deletion prohibited when implementing minGenome. Next, we used flux balance analysis on the published genome-scale metabolic model (GSM) for *Synechococcus* 2973 (37) to compute the maximum growth rate using 10 mmol/gDW hr CO_2_ (as basis), and default model parameters. minGenome was then implemented using this metabolic model while fixing the growth rate at the determined maximal value, and constraining all essential genes to be preserved. Longest contiguous genome regions to be deleted were thus iteratively identified. It should be noted that as long as the GSM is used to simulate carbon-limited phototrophic growth, the deletion regions identified here should be invariant of the specific model parameters used.

### Chemicals and Reagents

All enzymes were purchased from New England Biolabs (NEB, Ipswich, MA, USA) and ThermoFisher Scientific (Waltham, MA, USA). The molecular biology kits were obtained from Sigma-Aldrich (St. Louis, MO, USA). The chemicals, reagents and antibiotics used in this study were of analytical/HPLC grade and were procured from Sigma-Aldrich (St. Louis, MO, USA). The primers were ordered from Integrated DNA technologies (IDT, Coralville, IA, USA). The plasmids were sequenced by Genewiz® (South Plainfield, NJ, USA).

### Cultivation condition of the WT and mutants

The WT *Synechococcus* 2973 strain and mutant strains were cultivated and maintained in Caron chamber under 300μmoles.m^-2^.s^-1^ light and 0.5% CO_2_ at 38°C and rpm of 250 rpm. *E. coli* containing specific plasmids was cultivated overnight at 37°C in LB supplemented with appropriate antibiotic. Freezer stocks were maintained at -80°C in 7% DMSO for cyanobacterial strains and 25% glycerol for *E. coli* strains.

### Development of Cas3 mediated large random deletions

The engineered strains were constructed using the CRISPR-Cas3 editing technique novel to the cyanobacterial system. This is a multi-Cas system where Cas3 has the nuclease-helicase activity (26). In our previous study we developed a an inducible CRISPR plasmid pSL3577 (pRhRB-Cas3578), where the *cas* genes and the gRNA are controlled using the Rhamnose inducible promoter and Theophylline inducible riboswitch (Fig. S6) (27, 38).. The BsaI restriction site was used as the site for gRNA cloning in pSL3577 and this system does not require any predetermined repair template. The recombinant plasmid (pSL3578) was conjugated into the *Synechococcus* 2973 by triparental mating using a modified protocol. Briefly, the recombinant plasmids were transformed into the competent cells of WM6026 containing the pRL623 plasmid. This strain was used for conjugation. 2 mL of *E. coli* cells were grown overnight in SOB media supplemented with 30 μg/ml chloramphenicol and 250 mM DAP, which was diluted in 25 mL SOB and DAP (without antibiotic) and grown to an OD of 0.1-0.2. This 25 mL of exponentially grown conjugal strain was mixed with 5 mL of exponentially grown cyanobacterial culture and centrifuged at 2000 rpm. The mixed cells were washed gently and resuspended in 400 μL of BG-11 medium and evenly spread on to a HATF nitrocellulose filter paper (MilliporeSigma, St. Louis, MO, USA) that was placed on a BG-11 5% LB agar plate without antibiotic, prior to the experiment. The plates were incubated for 24 h under 100 μmoles.m^-2^.s^-1^ light and ambient air at 38°C to obtain mat growth of cyanobacterial cells. The filters were then carefully transferred to BG-11 agar plate with 50 μg/mL kanamycin and incubated under 300 μmoles.m^-2^.s^-1^ light and 0.05% CO_2_ until the lawn growth disappeared and individual colonies appeared. A transformant colony was selected randomly and grown in 20ml BG11+Kan media. Then it was induced for 24 h with 2g/L Rhamnose and 1mM Theophylline. Post induction, the culture was diluted and plated on antibiotic plate to obtain single colonies, which were tested for deletion using tiling PCR (26), whole genome sequencing (WGS) as reported earlier (27). Primers are listed in Table S7.

### Growth characteristics

The WT and the mutant strains were grown at 38 °C in 100 ml glass cultivation tubes of Multi-Cultivator photobioreactor (Photon Systems Instrument, Multi-Cultivator MC 1000, Czech Republic) containing 50 ml BG11 media under light intensity of 1500 (HL) μmoles.m^-2^.s^-1^. The aeration provided for growth was by bubbling either 1% CO_2_ mixed air or atmospheric air through the culture. Cells were inoculated at an OD720nm of 0.025-0.1 and were cultivated to an OD of 1-1.5. The growth rate and doubling time was calculated in the exponential phase of growth. All experiments were performed in biological replicates (at least n=3).

The absorption spectra of the WT and mutant strains were obtained at room temperature using Shimadzu UV-1800 spectrophotometer. The whole cell chlorophyll contents were calculated from the absorption spectra using the formulae obtained from Arnon et al., 1974 (39). The cultivated cyanobacterial cells were normalized on the basis of chlorophyll content to determine the quantum efficiency (40) of strains grown under growth condition using the FL-200 dual modulation PAM fluorometer with blue light activation as per previous protocol (41).

### Expressing sucrose transporter in the strain with minimized genome and sucrose measurement

The sucrose over-excreting strain was constructed as reported earlier (10, 27). Briefly, the cscB gene encoding for sucrose permease (source: *E. coli* (ATCC 700927)) was integrated at the neutral site 3 (NS3) of SG55 strain and streaked on Kanamycin plate for complete segregation. The sucrose productivity of SG55-cscB strain was compared with the highest sucrose producing strain of *Synechococcus* 2973 reported previously (10). The sucrose levels were determined following the previous protocol, briefly, cells were grown in 12 well-plate for a period of 3 days along with 1mM IPTG inducer added to the media. Post 3 days cells harvested (OD730nm reached ∼1) and centrifuged to obtain the supernatant which was used for sucrose measurement using the sucrose/D-glucose assay kit (Megazyme) (10, 27). The standard curve for sucrose and glucose were performed (27). Experiment was performed in biological (n=3) and technical (n=3) replicates.

## Acknowledgement

The authors are grateful to the funding agency for this study National Science Foundation (NSF) MCB-2037887 to HBP, NSF MCB-2037995 to GMC and NSF MCB-2037829 to CDM

## Authors Contribution

HBP, GMC and CDM conceived the study. AS, AB, MGS designed the experiment. AS and DL conducted the experiments. AS and AB analyzed the data. MSG performed the sequencing studies. DS and JIH performed the computational analysis. AS wrote the first draft of the manuscript. All the authors reviewed and revised the manuscript.

